# Information sharing for a coordination game in fluctuating environments

**DOI:** 10.1101/268268

**Authors:** Keith Paarporn, Ceyhun Eksin, Joshua S. Weitz

## Abstract

Collective action dilemmas pervade the social and biological sciences - from human decision-making to bacterial quorum sensing. In these scenarios, individuals sense cues from the environment to adopt a suitable phenotype or change in behavior. However, when cues include signals from other individuals, then the appropriate behavior of each individual is linked. Here, we develop a framework to quantify the influence of information sharing on individual behavior in the context of two player coordination games. In this framework, the environment stochastically switches between two states, and the state determines which one of two actions players must coordinate on. Given a stochastically switching environment, we then consider two versions of the game that differ in the way players acquire information. In the first model, players independently sense private environmental cues, but do not communicate with each other. We find there are two types of strategies that emerge as Nash equilibria and fitness maximizers - players prefer to commit to one particular action when private information is poor, or prefer to employ phenotypic plasticity when it is good. The second model adds an additional layer of communication, where players share social cues as well. When the quality of social information is high, we find the socially optimal strategy is a novel “majority logic” strategy that bases decision-making on social cues. Our game-theoretic approach offers a principled way of investigating the role of communication in group decision-making under uncertain conditions.

## 1. Introduction

In stochastically fluctuating environments, organisms adapt their behavior to increase their chances of survival. For instance, bacteria process information from extracellular cues to reduce their uncertainty about the environmental state and employ stochastic phenotype switching in proportion to the remaining uncertainty (Lopez et al., 2009; Perkins and Swain, 2009). However in many circumstances, coordination of behavior with others directly influences survival. In bacteria, some cooperative behaviors are facilitated by quorum sensing, a cell-to-cell communication system where individual cells secrete and sense autoinducer molecules to obtain information about the environment and to gauge local cell density (Miller et al., 2001; Henke and Bassler, 2004). Quorum sensing can induce formation of biofilms for protection against a host’s immune system, secretion of virulence factors to consolidate colonization of the host, motility, and many other behaviors (West et al., 2007; Nadell et al., 2008; de Kievit and Iglewski, 2000; Atkinson et al., 2006). In honeybee and ant colonies, individuals process and share information to collectively reach an informed collective decision about the best nesting site (Franks et al., 2002; Pratt et al., 2002). The individual-level mechanisms that produce collective behaviors in animal groups is an area of ongoing research (Couzin, 2009; Sasaki and Pratt, 2018). Inspired by these examples, the aim of this paper is to develop a game-theoretic framework in which to study the individual-level decision-making processes which produce collective behavior under environmental uncertainty and noisy communication.

There is an extensive body of work on individual (as opposed to group) decision-making in fluctuating environments (Perkins and Swain, 2009; Lopez et al., 2009). In many scenarios, an individual must match its phenotype or behavior to changing conditions by using sensory cues from the environment, signals from other individuals in the population, or both (Lachmann et al., 2000; Kussell and Leibler, 2005; Rivoire and Leibler, 2011; Donaldson-Matasci et al., 2010, 2013). An individual employing the optimal bet-hedging strategy diversifies behaviors at frequencies that mirror the posterior knowledge of environmental fluctuations. These results quantify an information-theoretic connection between the optimal population-level growth rate and the amount of information about the environment available to the individual. However, the resulting individual fitness is independent of the actions of others in the population. Hence, these models do not address the interplay between information and collective decision-making in fluctuating environments. In bacterial quorum sensing, the autoinducer signaling molecules that individual cells send serve as social cues that indicate local cell density (Miller et al., 2001; Nadell et al., 2008; Popat et al., 2015). Substantial experimental research has been done in recent years to unravel the mechanisms of this complex communication system. Nonetheless, questions remain regarding why such communication systems are utilized, particularly from an evolutionary standpoint (Whiteley et al., 2017).

Game theory offers a framework to explain rational behaviors when an individual’s well-being depends on the actions of others. To understand the role of communication in collective decisionmaking under uncertainty, we recognize there are two components to the decision-making process: 1) a *communication system*, or the way individuals acquire information and 2) *strategies*, or the way individuals use acquired information to make a decision. In this vein, two recent works have attempted to understand the role of communication in group coordination under uncertain fluctuating environments from a game-theoretic perspective. Pacheco et al. (2015) studies the evolutionary outcomes of communication systems in an *N*-person volunteer’s dilemma game, where all players adopt a majority rules strategy. Burgos and Polani (2016) consider how the choice of communication systems affect levels of cooperation between two populations of microbes in an information exchange game. In that model, each individual is assumed to follow a bet-hedging strategy. Both works assume the players’ strategies are fixed while the communication systems are evolvable, and do not investigate how players may adapt their strategies to a given communication system.

In this paper, we study a two-player, two-action game where the environment stochastically switches between two possible states. The state determines game payoffs, where the players must coordinate on the correct action corresponding to the environment. Here, we study two versions of this game in order to highlight the value of information sharing. In the first, players only receive a private cue from the environment. In the second, players receive a shared social cue in addition to private cues. By considering social interactions and their effects on a single stage payoff, our game-theoretic model differs from bet-hedging models that focus on the link between variation in individual strategies and long-term growth rates of the population. Additionally, complementary to Pacheco et al. (2015) and Burgos and Polani (2016), we seek to identify the optimal strategies that promote coordination given the limitations of a fixed, noisy communication system of variable fidelity. Most notable is a “majority logic” type strategy that is socially optimal when environmental sensing has intermediate reliability, and information sharing is very reliable. As we show, this allows an individual to act upon their inference of the environmental state when it is validated by social information.

## 2. Model and methods

### 2.1. A coordination game in fluctuating environments

We consider a two action, two player game that is played repeatedly in stages *t* = 1, 2,.… We denote the set of players 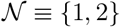 with generic member *i*, and the set of actions 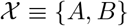. At each stage *t*, the environment *E*(*t*) takes one of two possible states, *e*_*A*_ or *e*_*B*_. If *E*(*t*) = *e*_*A*_ then *E*(*t* + 1) = *e*_*B*_ with probability *v*_*BA*_, and if *E*(*t*) = *e*_*B*_ then *E*(*t* + 1) = *e*_*A*_ with probability *v*_*AB*_. We assume that *E*(1) is arbitrarily determined and the switching probabilities *v*_*AB*_ > 0 and *v*_*BA*_ > 0 are fixed. Hence, the environmental state evolves according to a two-state Markov chain that spends a fraction 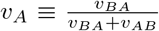 and 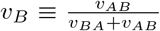 of time in states *e*_*A*_ and *e*_*B*_, respectively. Once *E*(*t*) is determined, players select their actions 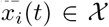. They do not know *E*(*t*) with certainty, and the state *E*(*t*) governs the players’ utility functions at each stage. We are interested in the average payoff of the players given the stochastically switching environment.

We define the game that is played at each stage as follows. Both must coordinate on action *A* (*B*) if *E*(*t*) = *e*_*A*_ (*e*_*B*_) to receive a payoff *b*_*A*_ > 0 (*b*_*B*_ > 0). If they mis-coordinate, or if they coordinate on the incorrect action, the payoff is zero. The environment-dependent payoff matrices are illustrated in Figure 1. Note that the payoff value to either player is the same. Thus, a generic player’s utility function is defined as

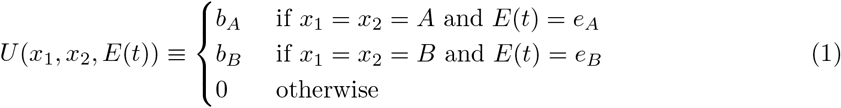

Given the realization of the environment *E*(*t*), a normal form game is played at stage *t* between the set of players 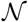 equipped with action set 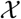, and with utility function *U*(·, ·, *E*(*t*)). We denote this normal form game by the triple 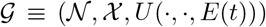, following standard game-theoretic notation. In the following, we introduce two models where players receive noisy information about *E*(*t*), and present our definition of fitness, which we refer to as *time-averaged payoffs*.

### 2.2. The game 𝒢_*p*_ with environmental sensing

At each stage *t*, suppose each player (*i* = 1, 2) independently senses the environment *E*(*t*) by receiving a private cue *α*_i_(*t*) ∈ {*e*_*A*_, *e*_*B*_}. The cue *α*_*i*_(*t*) matches the true environmental state with probability *p* ∈ [1/2,1], and mismatches with probability 1 – *p*.

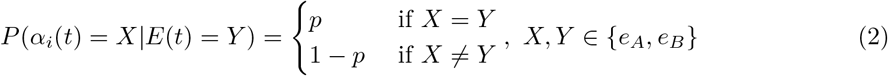

Thus, *α*_*i*_(*t*) is the output of a binary noisy channel of fidelity *p* whose input is *E*(*t*). We will refer to the parameter *p* as the sensing fidelity.

A *strategy* is a mapping from the set of cues {*e*_*A*_,*e*_*B*_} to the set of actions 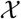. In other words, a strategy is a contingency plan or policy that a player adopts that assigns the action to take upon receiving a particular cue. Hence, it is a description of how a player makes informed decisions. Since there are two possible cues and two actions, each player can choose among 2^2^ = 4 strategies. We assume player *i* chooses only one strategy *s*_*i*_, which is fixed for all stages t = 1, 2,.… For notational convenience, we will denote a strategy by an ordered two-vector whose entries are either *A* or *B*. For instance, *s*_*i*_ = [*A*, *B*] denotes the strategy where *i* plays action *A* when *α*_*i*_(*t*) = *e*_*A*_ and action *B* when *α*_*i*_(*t*) = *e*_*B*_. We write *s*_*i*_(*e*_*A*_) = *A* and *S*_*i*_(*e*_*B*_) = *B*. We denote the set of all four strategies *S*^4^. When the context is clear, we will also omit the notation (*t*) indicating variables realized at stage *t*.

The list of all four strategies is given in Table 1. They are read “Only A” (OA), “Follow Cue” (FC), “FC bar” 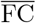
, and “Only B” (OB). When *p* = 1/2, *s*_*FC*_ and 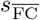 reduce to the strategy that uniformly randomizes between *A* and *B*.

**Figure 1:**
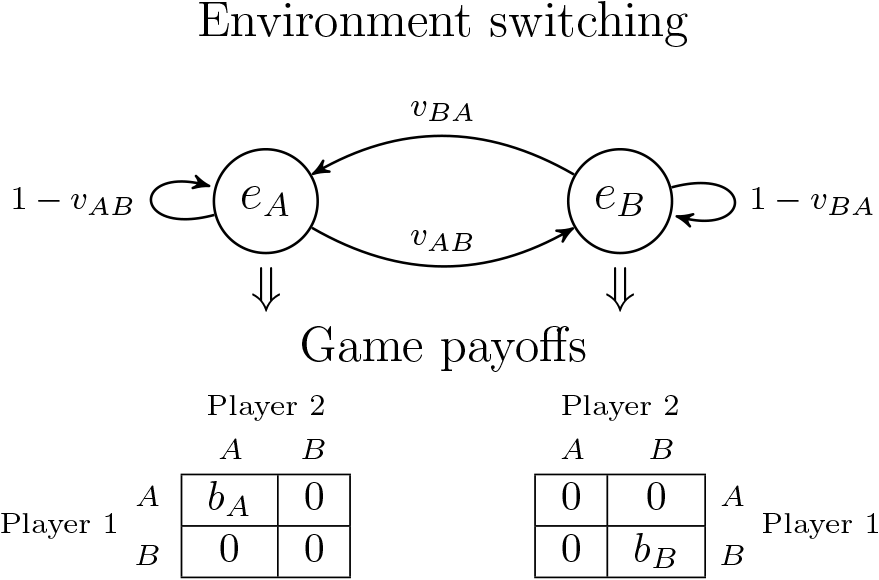
After each stage, the environment stochastically switches between *e*_*A*_ and *e*_*B*_ and determines the payoff matrix. In *e*_*A*_(*e*_*B*_), players need to coordinate on the *A*(*B*) action to receive a positive payoff *b*_*A*_(*b*_*B*_).

#### Remark 1.

These strategies, with the exception of 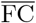, can be classified as a max likelihood estimate of the environmental state, depending on the environmental switching probabilities *v*_*A*_,*v*_*B*_ and private cue fidelity *p*. Specifically, the strategy

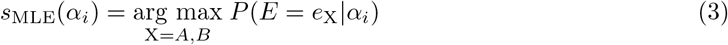

is precisely OA (*v*_*B*_ < *p* < *v*_*A*_), OB (*v*_*A*_ < *p* < *v*_*B*_), or FC (*v*_*A*_ < *p*, *v*_*B*_ < *p*). In this paper, we omit analysis of the *s*_MLE_ strategy because it simply corresponds to one strategy in Table 1 for any given set of parameters.

**Table 1:** List of all strategies in the game with no information sharing.

| $\alpha_i / s_i$ | $s_{\text{OA}}$ | $s_{\text{FC}}$ | $s_{\overline{\text{FC}}}$ | $s_{\text{OB}}$ |
| --- | --- | --- | --- | --- |
| $e_A$ | $A$ | $A$ | $B$ | $B$ |
| $e_B$ | $A$ | $B$ | $A$ | $B$ |

A measure of fitness is the fraction of time the players coordinate on the correct action, weighted by the benefits *b*_*A*_ and *b*_*B*_ accordingly. We calculate this measure as follows. Consider the environment *E*(*t*), which evolves according to the two-state Markov chain with stationary distribution (*v*_*A*_, *v*_*B*_). Additionally, the players’ cues are drawn independently of each other, but conditionally on the state *E*(*t*). Since neither the cues nor player actions affect the environment switching probabilities, the fraction of time spent in the aggregate state (*α*_1_, *α*_2_, *E*) is given by the following stationary distribution

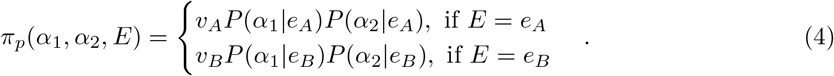

For instance, the value of the stationary distribution π_*p*_ at the entry (*e*_*A*_, *e*_*A*_, *e*_*B*_) is *v*_*B*_(1 – *p*)^2^. We define the *time-averaged payoff* as the expected utility with respect to the stationary distribution,

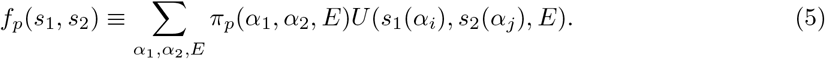

Via (5), we define the normal form game 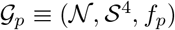 played between players 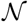 equipped with strategy space *S*^4^, and with utility function *f*_*p*_ for a given sensing fidelity *p* ∈ [1/2,1]. Figure 2 shows a diagram of the stage game and information system that underlies 𝒢_*p*_. We can represent 𝒢_*p*_ with the following 4 × 4 matrix of time-averaged payoffs *f*_*p*_(*s*_1_, *s*_2_), 
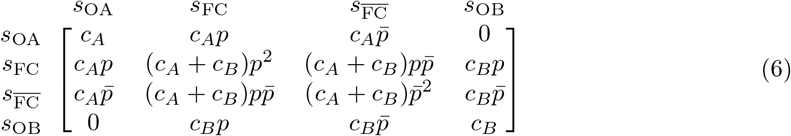

where 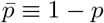 and 
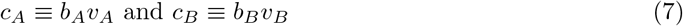
are the relative benefits of each environmental state. Note the payoff matrix is symmetric, which gives 10 unique strategy profiles. Therefore, the identity of the players do not matter. An expression to calculate each entry of (6) is given in Appendix A of the SI. We note that a “normalized” fitness value for all the entries above can be attained by dividing by *c*_*B*_. This captures all relative payoffs in terms of a single environment-related parameter, 
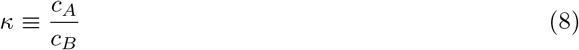
We call *κ* the *ratio of relative benefits*, and it is positive and nonzero. We will later see in Section 3 that *κ* is useful for parameterizing Nash equilibrium and fitness maximizer regions.

**Figure 2:**
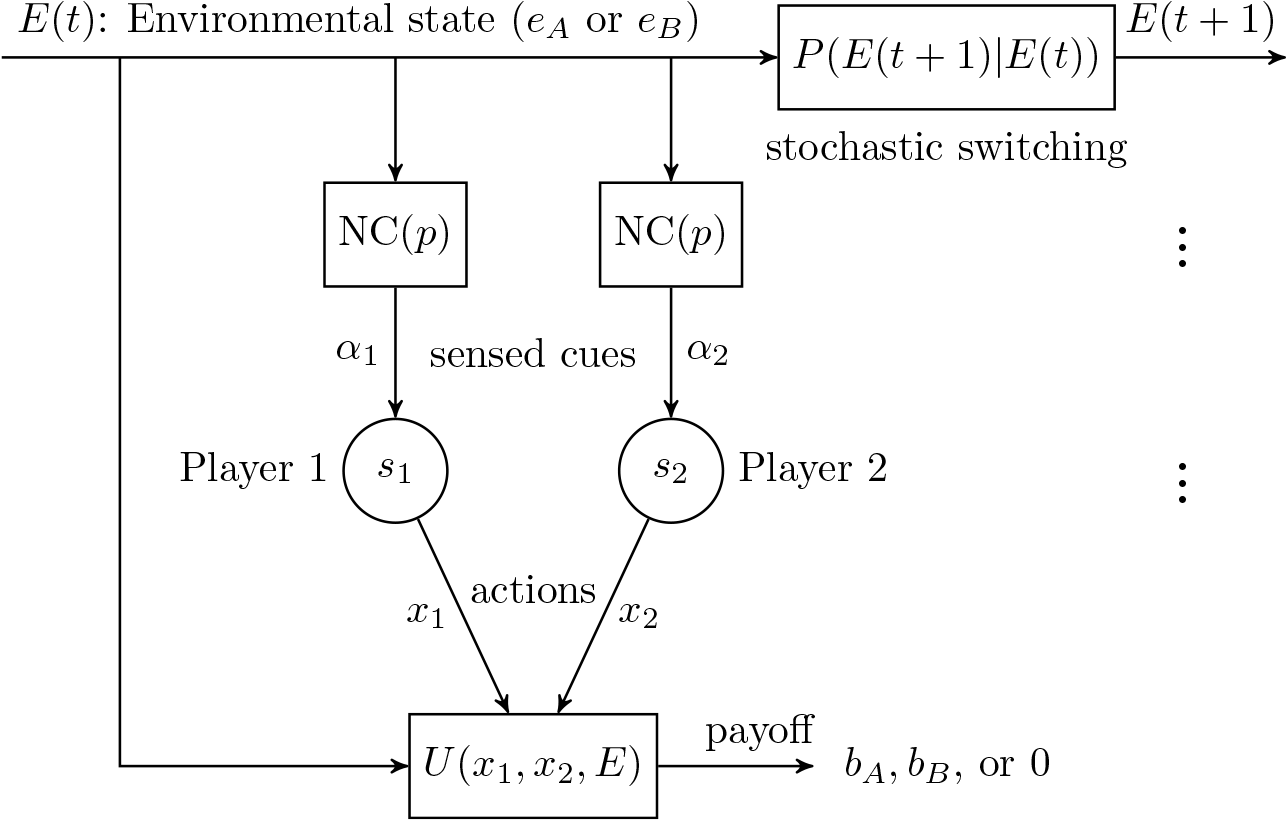
Diagram of the game with no information sharing 𝒢*p*. At stage *t*, each player independently senses a cue *α*_*i*_ directly from the environment *E*(*t*) through a binary noisy channel with fidelity *p*, denoted here as NC(*p*). That is, *α*_*i*_ = *E*(*t*) with probability *p*, and *α*_*i*_ ≠ *E*(*t*) with probability 1 – *p*. Player i’s strategy *s*_*i*_(*α*_*i*_) determines its action *x*_*i*_ ∈ {*A*,*B*}. The payoff *U*(*x*_1_,*X*_2_,*E*) (eq. (1)) to both players is either *b*_*A*_, *b*_*B*_, or 0, which is determined by the current actions of both players and the current environment. The environmental state stochastically switches to *E*(*t* + 1) at stage *t* + 1.

#### Remark 2.

The average payoff (5) can also be viewed as the *ex-ante* expected utility in the one-shot Bayesian game *B*_*p*_ consisting of the set of players 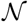, action set *A*, external states {*e*_*A*_, *e*_*B*_}, type space {*e*_*A*_, *e*_*B*_}, utility *U* (eq. (1)), beliefs *P*(*α*_1_,*α*_2_,*E*), and a common prior on *E*, *P*(*e*_*A*_) = *v*_*A*_ and *P*(*e*_*B*_) = *v*_*B*_. The ex-ante expected utility is defined as the expectation of *U* with respect to the belief *P*(*α*_1_, *α*_2_, *E*), which is the induced probability distribution over the aggregate state of the world and coincides with (4). In an ex-ante setup, players evaluate their utilities before receiving their signal *α*_*i*_, and therefore must reason about the possible environmental states, the signal of the other player, and its own signal. The (normal form) Nash equilibrium solution concept applied to the game 𝒢_*p*_ coincides with the definition of ex-ante Bayesian Nash equilibrium (BNE) of *B*_*p*_. A Bayesian Nash equilibrium describes a state of rationality where no player can profitably deviate by changing its strategy given its belief of the world. We focus our attention on the normal-form formulation 𝒢_*p*_ in this paper, keeping in mind that a Nash equilibrium in 𝒢_*p*_ can also be interpreted as a Bayesian Nash equilibrium of *B*_*p*_. A treatment of Bayesian games can be found in Ch. 6 of Vega-Redondo (2003).

### 2.3. The game 𝒢_*pq*_ with information sharing

The game 𝒢_*p*_ is extended by allowing players to share their private cues with each other before deciding on an action. At stage *t*, after players sense *α*_*i*_(*t*), player *i* sends a social cue *β*_*j*_(*t*) to player *j* (*j* ≠ *i*), which matches *i*’s private cue *α*_*i*_(*t*) with probability *q* ∈ [1/2,1].

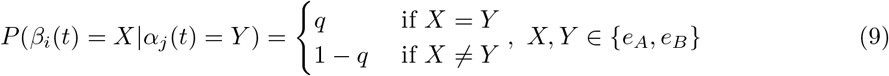

Thus, *β*_*i*_(*t*) is the output of a binary noisy channel of fidelity *q* whose input is *α*_*j*_ (*t*). We will refer to parameter *q* as the *sharing fidelity*. We assume here that players signal honestly, i.e. each player attempts to share their true private cue. This is a reasonable assumption because the players’ interests are aligned - they must attempt to coordinate. In different contexts where player interests conflict, dishonest signalling becomes a rational alternative. For example, male fiddler crabs with inferior claws can bluff fighting ability to ward off other males (Backwell et al., 2000).

Player *i*’s information is now composed of the pair *y*_*i*_(*t*) ≡ (*α*_*i*_(*t*), *β*_*i*_(*t*)) ∈ {*e*_*A*_,*e*_*B*_}^2^. Similarly for this model, a strategy is a mapping from the set of information pairs {*e*_*A*_, *e*_*B*_}^2^ to actions, so player *i* can now choose among 2^4^ = 16 strategies. We represent strategies as ordered four-vectors whose entries are *A* or *B*. For instance, *s*_*i*_ = [*A*, *B*, *B*, *A*] is the strategy where *i* plays action *A* when *y*_*i*_ = (*e*_*A*_, *e*_*A*_), *B* when *y*_*i*_ = (*e*_*A*_, *e*_*B*_), *B* when *y*_*i*_ = (*e*_*B*_, *e*_*A*_), and *A* when *y*_*i*_ = (*e*_*B*_, *e*_*B*_). We note that the four strategies available in 𝒢_*p*_ are also strategies in this game. They are now represented by the vectors *s*_OA_ = [*A*, *A*, *A*, *A*], *s*_FC_ = [*A*, *A*, *B*,*B*], 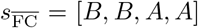, *s*_OB_ = [*B*, *B*, *B*, *B*]. These four strategies base decision-making either on no information at all (OA and OB), or only on the private cue *α*_*i*_ (FC and 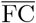). The 12 new strategies base decisions on both private and shared signals. The set of all 16 strategies is denoted *S*^16^.

Similarly to (4), we calculate the time-averaged payoff by considering the aggregate state (*y*_1_, *y*_2_, *E*), which occurs a fraction of the time according to the following stationary distribution 
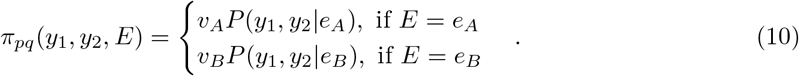
One can expand *P*(*y*_1_,*y*_2_|*E*) = *P*(*α*_1_|*E*)*P*(*α*_2_|*E*)*P*(*β*_1_|*α*_2_)*p*(*β*_2_|*α*_1_). For instance, the value of the stationary distribution at the entry ((*e*_*A*_, *e*_*A*_), (*e*_*A*_, *e*_*B*_), *e*_*A*_) is *v*_*A*_*p*^2^*q*(1 – *q*). The *time-averaged payoff* at (*p*, *q*) ∈ [1/2, 1]^2^ is defined as

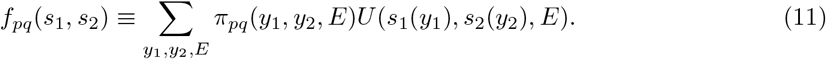

Via (11) and following the arguments from Remark 2, we define the normal form game 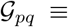 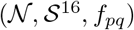 played between the set of players 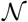 equipped with strategy space *S*^16^, and with utility function *f*_*pq*_ for a given pair of sensing and sharing fidelities (*p*, *q*) ∈ [1/2,1]^2^. Figure 3 shows a diagram of the stage game and communication system that underlies 𝒢_*pq*_. An expression to calculate each entry of the resulting 16 × 16 symmetric payoff matrix is given in Appendix A of the SI. Given the symmetry of the payoff matrix, there are 136 unique strategy profiles. A summary of all relevant parameters that define the games 𝒢_*p*_ and 𝒢_*pq*_ is listed in Table 2.

**Figure 3:**
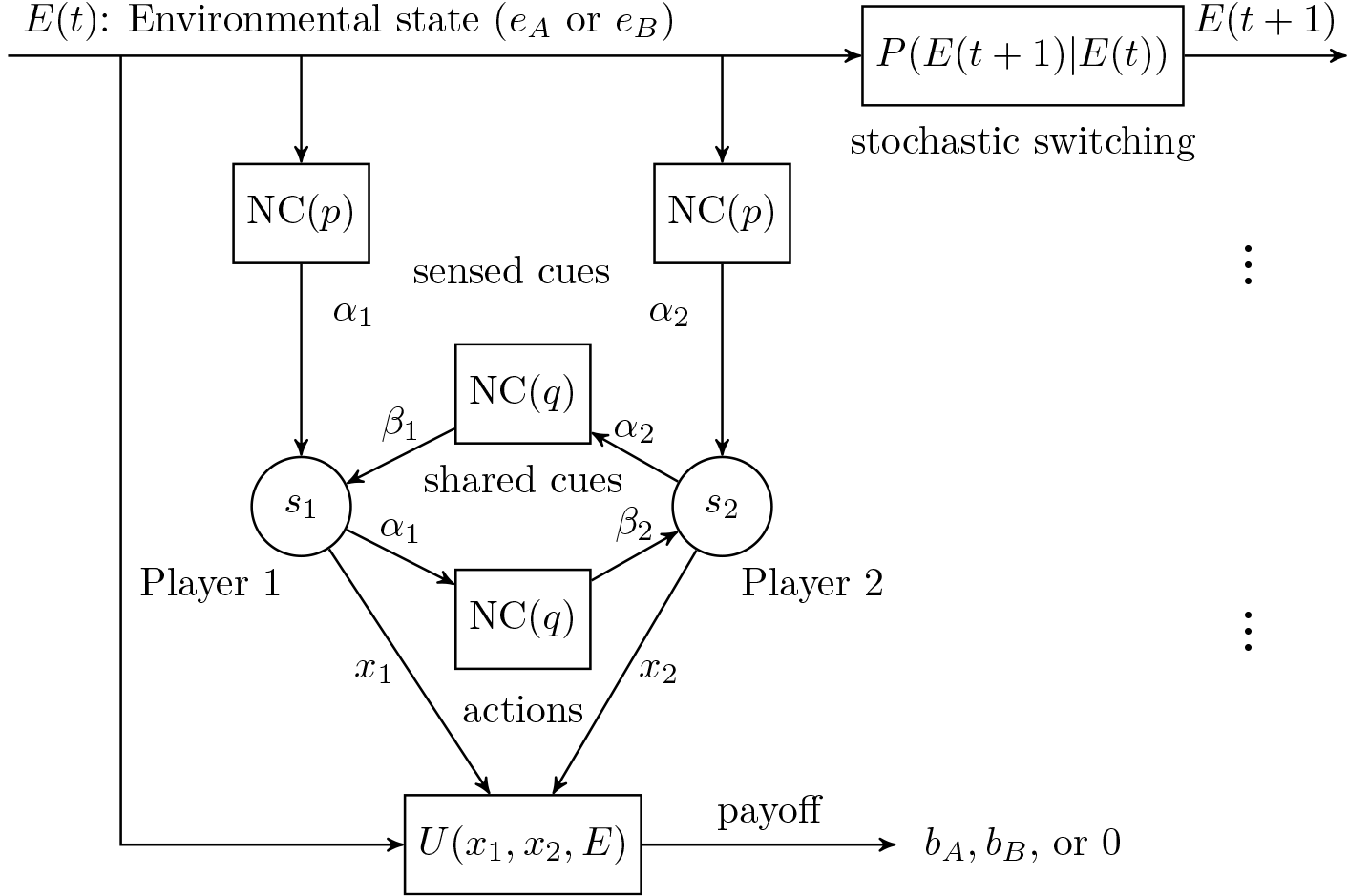
Diagram of the game with information sharing 𝒢_*pq*_. At stage *t*, each player independently senses a private cue *α*_*i*_ from the environment *E*(*t*) through a binary noisy channel with fidelity *p*, denoted here as NC(*p*). Then, each player receives a signal *β*_*i*_ from the other’s private cue through a separate noisy channel of fidelity *q*. Given player *i*’s information (*α*_*i*_,*β*_*i*_), its action is determined by its strategy, *s*_*i*_(*α*_*i*_,*α*_*i*_) = *X*_*i*_. The payoff *U*(*x*_i_,*X*_2_,*E*) (eq. (1)) to both players is either *b*_*A*_, *b*_*B*_ or 0, which is determined by the actions of both players and the current environment. The environmental state stochastically switches to *E*(*t* + 1) at stage *t* + 1.

**Table 2.**
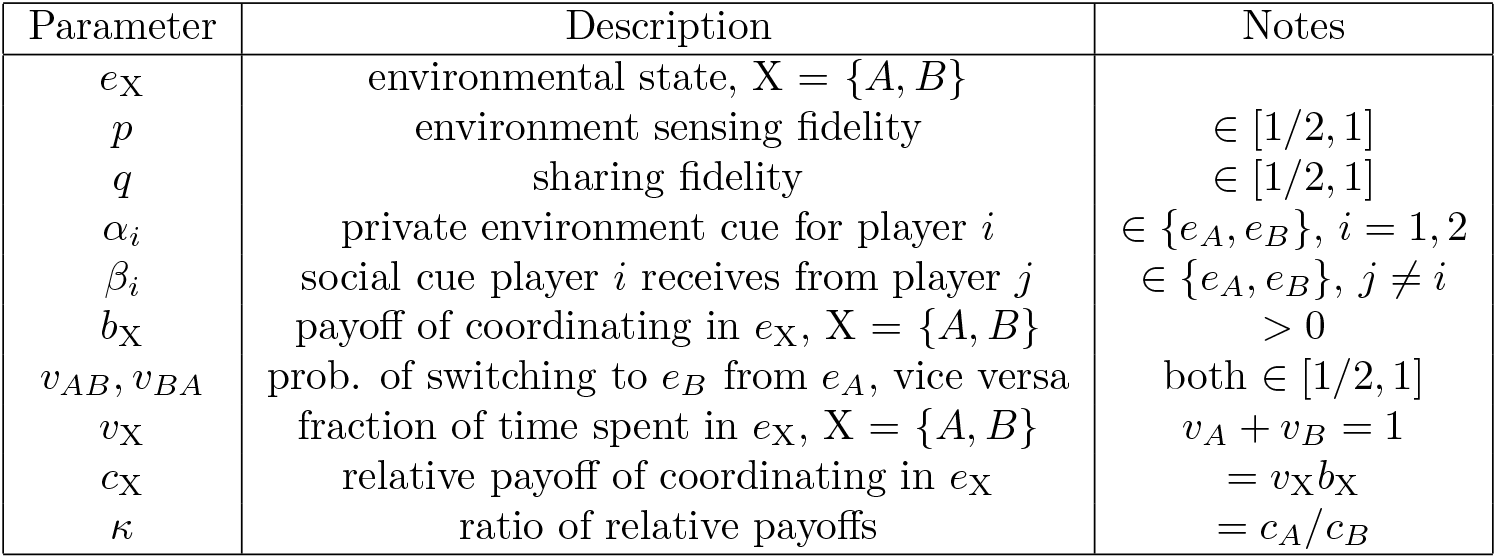
List of parameters that define the games 𝒢_*p*_ and 𝒢_*pq*_.

#### Remark 3.

We note that the payoff matrices of 𝒢_*p*_ and 𝒢_*pq*_ are symmetric. Hence, the identity of the player (player 1 or 2) does not matter. Such games fall into the class of potential games (Monderer and Shapley, 1996; Sandholm, 2001). A game is a potential game if the incentives of the players align with a global *potential* function. Here, the change in payoff from a unilateral deviation of a single player is equivalent to the change in global potential, given all other players remain the same. In our formulation, the potential function is the average player fitness, or welfare.

## 3. Results

We consider the Nash equilibrium solution concept and fitness maximizing strategies as notions of rational and optimal behavior, respectively. The comparison of the fitness maximizing strategies between the two models (𝒢_*p*_ and 𝒢_*pq*_) presented offers a principled way to investigate the role of sensing and communication in collective decision-making in systems such as quorum sensing.

Specifically, we are interested in how the quality of the communication system, defined by the parameter *p* for 𝒢_*p*_ and (*p*, *q*) for 𝒢_*pq*_, dictates which strategy profiles are Nash equilibria and fitness maximizers. A Nash equilibrium is the classical solution concept in game theory which describes a rational steady-state configuration where no player has an incentive to deviate from its strategy. A Nash equilibrium in 𝒢_*p*_ is a strategy profile (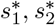) that satisfies

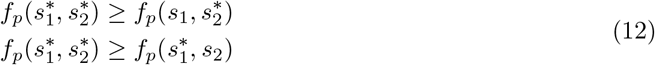

for all *s*_1_ ∈ *S*^4^, 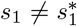, and for all *s*_2_ ∈ *S*^4^, 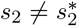. A strict Nash equilibrium is a Nash equilibrium satisfying (12) with strict inequality. We say a strategy profile (ŝ_1_, ŝ_2_) is a *fitness maximizer* at *p* if it satisfies

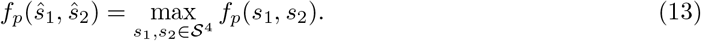

The same definitions above apply for 𝒢_*pq*_, where *f*_*p*_ is replaced by *f*_*pq*_, and *S*^4^ by *S*^16^. Because the payoff matrices of 𝒢_*p*_ and 𝒢_*pq*_ are symmetric, the following fact holds: in 𝒢_*p*_ and 𝒢_*pq*_, the fitness maximizer(s) is necessarily a Nash equilibrium. However, the converse is not true. We note that in our model, there is no fitness cost to having higher sensing and sharing fidelities *p* and *q*. Our aim is not to investigate such evolutionary tradeoffs, but to identify the types of strategies that ensure coordination in fluctuating environments given that the players utilize a communication system of quality (*p*, *q*) ∈ [1/2,1]^2^.

### 3.1. Nash equilibria and fitness maximizers in 𝒢_*p*_

The base game 𝒢 (see Eq. (1)) admits a single strict Nash equilibrium at the correct coordinated action. We might expect the game 𝒢_*p*_, represented by (6), also has a coordination structure. That is, the players are better off if they coordinate on the same strategy in *S*^4^. In other words, they should play the same action given they receive the same signal. We can show that this intuition is indeed correct and state the following fact: For *p* ∈ [1/2,1], all Nash equilibria and fitness maximizers of 𝒢_*p*_ are necessarily symmetric strategy profiles, 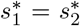. The proof of this fact, given in Appendix B of the SI, relies on showing that for any asymmetric strategy profile, there is one player that can switch to the other player’s strategy to improve the fitness *f*_*p*_.

Due to this result, we can limit our analysis to the four symmetric strategy profiles. To refer to symmetric strategy profiles (*s*, *s*) for the four strategies we simply write “OA” to denote (*s*_OA_,*s*_OA_), and similarly for the other three. Their resulting fitnesses correspond to the diagonal entries of (6).

We find that OA and OB are always Nash equilibria regardless of the value for sensing fidelity *p* and the relative benefits *c*_*A*_, *c*_*B*_. We find that 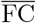 is not a Nash equilibrium for any set of parameters. Figure 4 (left) illustrates the Nash equilibrium region of FC. We can characterize the equilibrium region of FC in the game 𝒢_*p*_ with respect to just two parameters, (*p*, *κ*) ∈ [1/2,1] × (0, ∞), where we recall *κ* = *c*_*A*_/*c*_*B*_ is the ratio of relative benefits. This is possible by normalizing the fitness, *f*_*p*_/*c*_*B*_, and applying (12) to solve for the region’s conditions on *p* and *κ*.

Figure 4 (right) illustrates the regions where OA, OB, and FC are fitness maximizers. While OA and OB are Nash equilibria everywhere, there is a unique fitness maximizer (OA,OB, or FC), except for on the boundaries dividing each region, for any given (*p*, *κ*) value in [1/2,1] × (0, ∞). On the boundary lines, the fitnesses of the strategy profiles that are separated coincide. We note here that the region where FC maximizes fitness is a subset of its Nash equilibrium region. This reveals regions in the parameter space where FC is a suboptimal Nash equilibrium to the OA/OB strategies.

**Figure 4:w.**
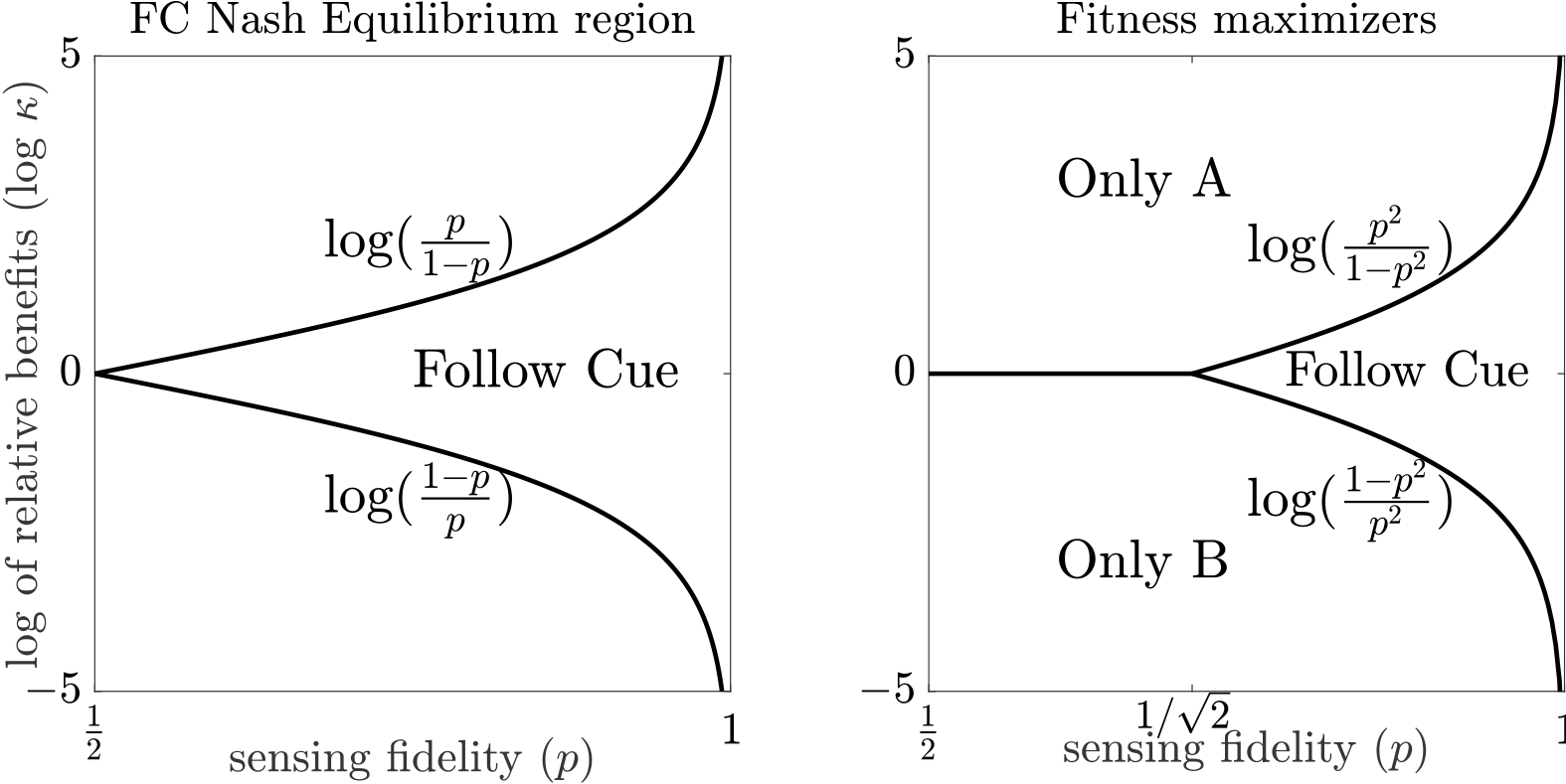
Characterization of strategies in the game 𝒢_*p*_ with respect to the parameter space (*p*, *κ*) with 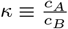. ranging in log scale from -5 to 5. Log scale is used to show symmetry between the ranges 0 < *κ* < 1 and 1 < *κ* < ∞. The quantity *κ* is the ratio of relative benefits between environment *e*_*A*_ and *e*_*B*_. (Left) The region where FC is a strict Nash equilibrium: 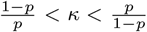. The strategies OA and OB are Nash equilibria everywhere. (Right) Disjoint regions where the strategies OA, OB, and FC are fitness maximizers. 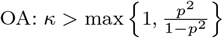. 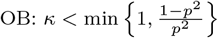. FC: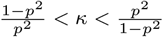

The fitness for FC increases quadratically in *p*: *f*_*p*_(FC) = (*c*_*A*_ + *c*_*B*_)*p*^2^. Hence, with higher sensing fidelity, players are better able to infer the correct environmental state and select the correct action. The *p*^2^ term appears because each player is independently sensing *E*, and coordination depends on both players independently receiving the correct cue which occurs with probability *p*^2^. The value of employing FC diminishes as *κ* → ∞ or 0. In either extreme, one environment is favored over the other. Either one occurs far more frequently over the long run, or its coordination benefit outweighs that of the other state. Hence, players do best by adopting either OA or OB (corresponding to the environment that is favored) instead of FC in these situations. FC is most desirable when *κ* ≈ 1, where either the environment fluctuates frequently and *b*_*A*_ ≈ *b*_*B*_, or a rare environment offers an enormous coordination benefit compared to the other. These are situations where acting on knowledge of the environment is most crucial. By playing OA or OB, players miss out on half of the fitness benefit opportunity whereas players employing FC are able to adapt to changing conditions.

The *s*_FC_ strategy possesses similarities to the optimal individual bet-hedging strategy that maximizes long-term population growth rate in fluctuating environments (Kussell and Leibler, 2005; Donaldson-Matasci et al., 2008, 2010) because it adapts actions based on the player’s inference on the environmental state. However in our model, the social context of coordination renders it suboptimal when private information is unreliable (low *p*), though it is optimal for sufficiently high values of *p*.

### 3.2. Nash equilibria and fitness maximizers in 𝒢_*pq*_

We now turn to the game 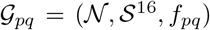. The Nash condition remains the same as in (12), except the time-averaged payoff is now given by *f*_*pq*_ (11). In comparison to 𝒢_*p*_, the parameter space of the information sharing game 𝒢_*pq*_ has another dimension, the social fidelity *q*. We search for the Nash equilibrium strategy profiles and fitness maximizers of 𝒢_*pq*_ over the parameter space (*p*, *q*, *κ*) ∈ [1/2,1]^2^ × (0, ∞). First, we observe that unlike 𝒢_*p*_, the coordination structure in the new set of strategies *S*^16^ is not preserved in 𝒢_*pq*_. Indeed, we find that the asymmetric strategy profile (*s*_1_,*s*_2_) with *s*_1_ = *s*_FC_ = [*A*, *A*, *B*, *B*] and *s*_2_ = [*A*, *B*, *A*, *B*] is a Nash equilibrium in a region of low sensing fidelity *p* and high sharing *q* (see SI Appendix C for derivation). Hence, not all Nash equilibria are necessarily symmetric strategy profiles for all (*p*, *q*) ∈ [1/2,1]^2^, and we cannot restrict our search for Nash equilibria and fitness maximizers to symmetric strategy profiles. In essence, players may now select new strategies that utilize the social cues as a means to coordinate.

There are 136 unique strategy profiles (by symmetry). In Figure 5, we display the multiplicity of Nash equilibria across the range of fidelity parameters *p* and *q*. Here, we do not count OA and OB because they are always Nash equilibria. We find the maximum number of Nash equilibria in the upper left region of the parameter space (*p*, *q*) ∈ [1/2,1]^2^, where private information is unreliable but information sharing has high fidelity. This suggests reliable social cues are utilized as a means to coordinate (see Section 4 for further discussion).

The strategy profiles that maximize fitness are shown in Figure 6 in the parameter space (*p*, *q*) ∈ [1/2, 1]^2^ and for cross-sections of *κ*. We note that the set of fitness maximizers consist only of symmetric strategy profiles. We find that OA, OB, and FC are fitness maximizers, along with a new type of strategy we term “Majority Logic” (ML). When environment *A* is favored (*κ* > 1), ML_*A*_ appears as a fitness maximizer, and similarly ML_*B*_ when *κ* < 1. These strategies are written

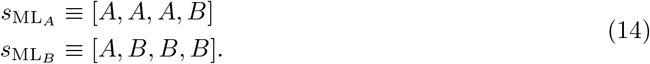
Recall that in our strategy vector notation, [*A*, *A*, *A*, *B*] is the strategy where the player chooses action *A* when *y*_*i*_ = (*e*_*A*_, *e*_*A*_), action *A* when *y*_*i*_ = (*e*_*A*_, *e*_*B*_), action *A* when *y*_*i*_ = (*e*_*B*_, *e*_*A*_), and action *B* when *y*_*i*_ = (*e*_*B*_, *e*_*B*_). The majority logic strategies ML_*A*_ and ML_*B*_ were not previously available in 𝒢_*p*_. Their fitnesses increase quadratically in *q*, and their functional forms are given in Appendix A.

The notation ML_*A*_,ML_*B*_ differentiates which action is assigned, *A* or *B*, to the cues *y*_*i*_ = (*e*_*A*_, *e*_*B*_) and (*e*_*B*_, *e*_*A*_), i.e. the middle two entries in the strategy vector. We will write just “ML” when generally speaking of the majority logic strategies (e.g. when *κ* = 1).

The region where the ML strategies maximize fitness is characterized by intermediate sensing fidelity *p* and high sharing fidelity *q*. The shared cues *β*_*i*_ are highly accurate so each player will have reliable knowledge of the other’s private cue. Our interpretation of why ML thrives in this regime is as follows. With reliable information sharing, players can detect when their private cues agree and when they disagree. As an example, consider when both players employ strategy *s*_mL_*A*__ = [*A*, *A*, *A*, *B*] and suppose environment *A* is favored over *B* (*κ* > 1). When both of their private cues coincide, i.e. *y*_1_ = *y*_2_ = (*e*_*A*_,*e*_*A*_) or (*e*_*B*_, *e*_*B*_), they choose the correct action. This occurs with overall probability *p*^2^*q*^2^. When they believe their private cues disagree with each other (e.g. *y*_1_ = (*e*_*A*_,*e*_*B*_) and *y*_2_ = (*e*_*B*_, *e*_*A*_)), they will both play action *A*, and obtain a positive payoff with overall probability *v*_*A*_*P*(1 – *p*)*q*^2^. Moreover, the above situations will occur far more frequently than players obtaining polarized beliefs (e.g. *y*_1_ = (*e*_*A*_, *e*_*A*_) and *y*_2_ = (*e*_*A*_, *e*_*A*_)), where they mis-coordinate actions with probability *p*(1 – *p*)(1 – *q*)^2^. The ML_*A*_ or ML_*B*_ strategy profile appears as the fitness maximizer depending on which environment is favored (Figure 6 right).

The *s*_ML_*A*__ strategy vector [*A*, *A*, *A*, *B*] ([*A*, *B*,*B*,*B*]) differs from *s*_FC_ = [*A*, *A*, *B*,*B*] and *s*_OA_ = [*A*, *A*, *A*, *A*] by only one action, whereas s_FC_ differs from *s*_OA_ by two actions (similarly for *s*_ML_*B*__ and *s*_OB_). Thus, we interpret *s*_mL_ to be a hybrid of *s*_FC_ and *s*_OA_. It acts as an estimator of the environment when the posterior belief on the environmental state is very high. This is the reasoning for the term “Majority Logic” - players act upon their inference of the environmental state only when their private information is validated by social information, i.e. when *y*_*i*_ = (*e*_*A*_, *e*_*A*_) or (*e*_*B*_, *e*_*B*_). Otherwise, when *y*_*i*_ = (*e*_*A*_,*e*_*B*_) or (*e*_*B*_,*e*_*A*_), the posterior is not as high and the player disregards its information altogether, playing the default action *A*.

**Figure 5:**
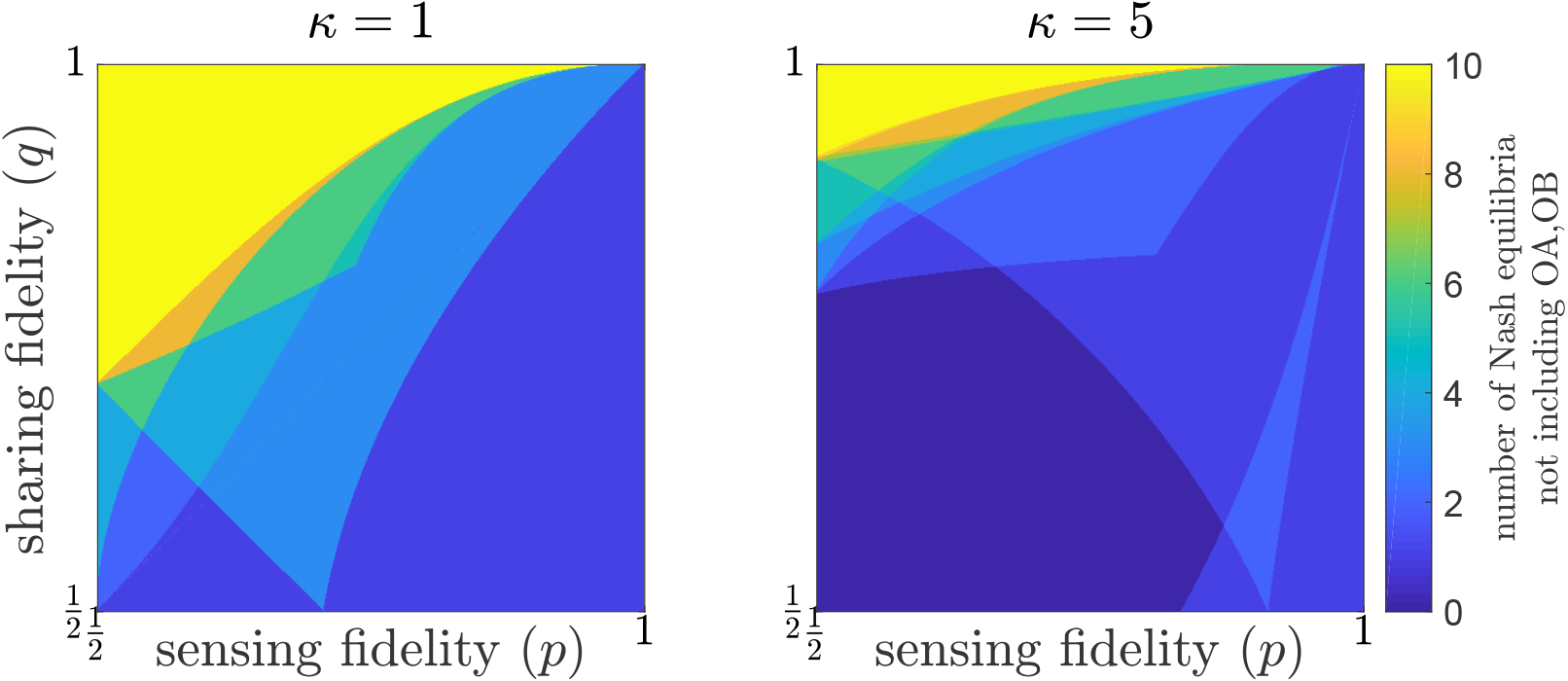
The number of Nash equilibria not including OA or OB, that exist across fidelity values (*p*, *q*) ∈ [1/2,1]^2^, for *κ* = 1 (Left) and *κ* = 5 (Right) in the game 𝒢_*pq*_. The data is numerically calculated by sweeping through (*p*, *q*) ∈ [1/2,1]^2^ in a uniform grid of spacing .001, and exhaustively verifying whether each 136 strategy profile satisfies its Nash condition, given in (12). We verify whether a strategy profile is a Nash equilibrium or not by using a simplified expression of the Nash condition in (12) that is amenable to numerical evaluation, given in Appendix C of the SI.

**Figure 6:**
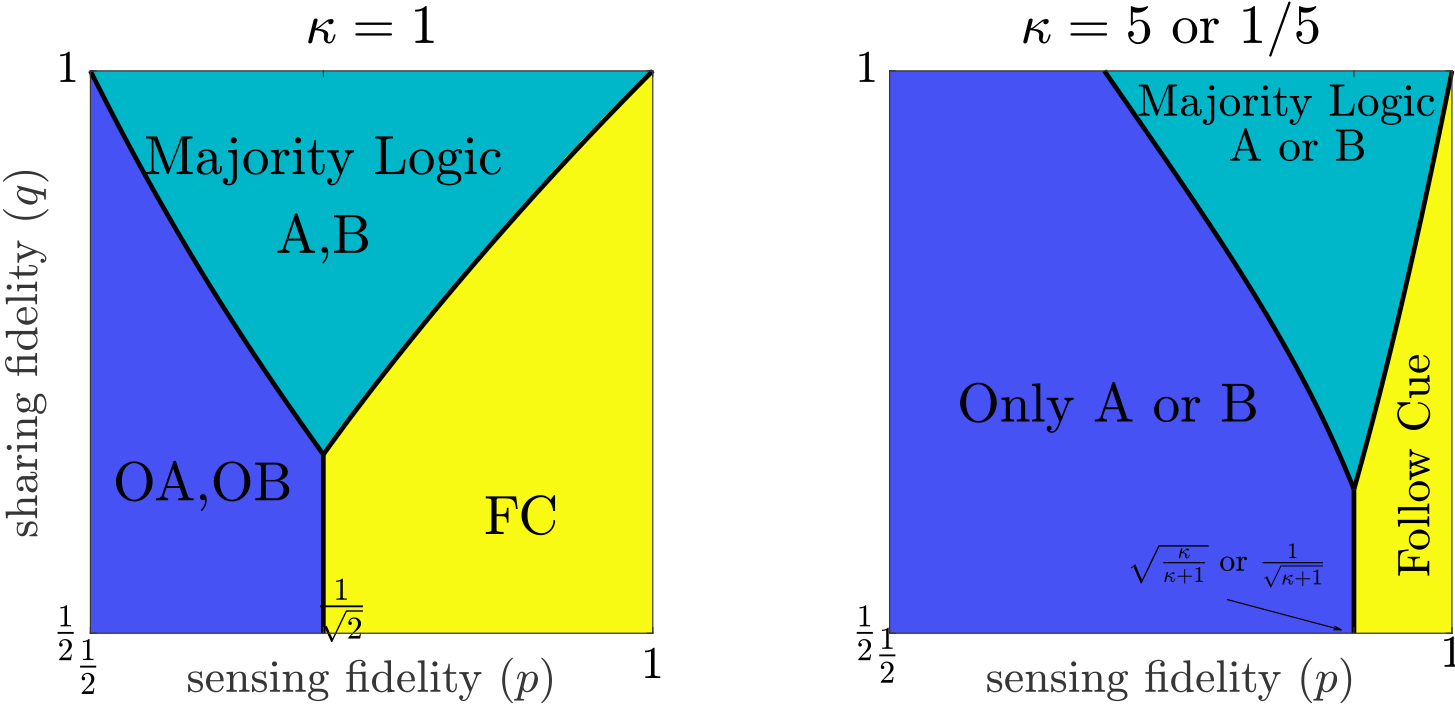
Three classes of strategies emerge as the fitness maximizers in 𝒢_*pq*_: 1) the pure strategies OA and OB, 2) follow cue (FC), and 3) majority logic (ML_*A*_ and ML_*B*_). The regions are drawn by first sweeping (*p*, *q*) ∈ [1/2,1]^2^ in a uniform grid with spacing .001, and exhaustively searching all 136 strategy profiles for the fitness maximizer. Each grid cell is then filled with the unique color corresponding to the fitness-maximizing strategy profile. Upon observing the emergence of the three strategy classes, the boundaries are analytically solved by equating their fitnesses. Hence, results are accurate within a spacing tolerance of .001. (Left) When *κ* = 1, players do not prefer any environment over the other. Hence, OA and OB give the same fitness value and we indicate this by OA/OB (similarly for ML_*A*_/ML_*B*_). (Right) When *κ* ≠ 1, the region boundaries are the same for both 5 and 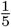 but the fitness maximizer is either OA or OB, depending on whether *κ* is greater or less than 1, respectively (similarly, either ML_*A*_ or ML_*B*_).

### 3.3. The fitness value of information sharing

Majority logic stands alone among all possible strategies in 𝒢_*pq*_ that can outperform the fitness maximizers of 𝒢_*p*_ (OA, OB, and FC). However, recall from Figure 6 (Left, for example) that it requires the sharing fidelity *q* to be sufficiently high for a fixed sensing fidelity *p*. This critical threshold value, which we denote with *q*_*c*_(*p*), is the value of *q* above which the majority logic strategy is the fitness maximizer. Hence, the values of *q*_*c*_(*p*) are parameterized by the boundary lines that separate the ML strategies from OA/OB and FC in Figure 6. Full parameterizations of *q*_*c*_(*p*) are given in Appendix D of the SI.

The critical thresholds *q*_*c*_(*p*) when the ML strategies outperform the optimal strategies of 𝒢_*p*_ suggests there is no value for players to share signals unless sharing fidelity is sufficiently high, *q* > *q*_*c*_(*p*). Higher sharing fidelity is needed for extreme values of *p*: *q*_*c*_(*p*) increases up to 1 as *p* decreases towards 1/2, as well as when *p* increases towards 1. When *p* is near 1, players prefer FC because they are able to independently detect the correct environment with very high probability and act accordingly. When *p* is near 1/2, private cues are effectively random because they contain no information about *E*. Consequently, the shared signals are also effectively random. Players then receive any of the four signals *y*_*i*_ with equal probability. By employing ML in this regime, players will miscoordinate more often than they would if they committed to OA or OB.

We are also interested in quantifying the fitness benefit of sharing signals over no sharing. Let 
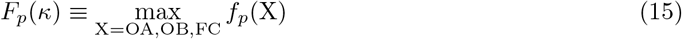

be the maximum fitness attainable in the game 𝒢_*p*_ at (*p*, *κ*), and let 
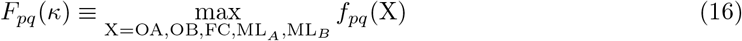
be the maximum fitness attainable in the game 𝒢_*pq*_ at (*p*, *q*, *κ*). Then we define the *fitness value of information sharing* as 
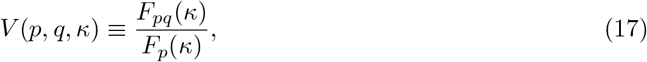
which gives the ratio of the maximum fitness in 𝒢_*pq*_ to the maximum fitness in 𝒢_*p*_ at the parameters (*p*, *q*, *κ*). By definition, *V*(*p*, *q*, *κ*) = 1 for *q* ≤ *q*_*c*_(*p*). In other words, there is no fitness benefit to sharing signals when *q* does not exceed the threshold *q*_*c*_(*p*). A contour map of *V* is shown in Figure 7. When *κ* =1 (Figure 7, Left), *V* is maximized at 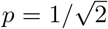 and *q* =1, giving 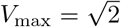 (using Eq. (A.12) and divide by *c*_*B*_). Thus, information sharing can improve fitness by approximately 41% when the environment fluctuates frequently. For *κ* > 1, it is maximized at *q* = 1, 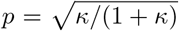 (Figure 7, Right). The general form of *V*_max_(*κ*) is the following piecewise continuous function, 
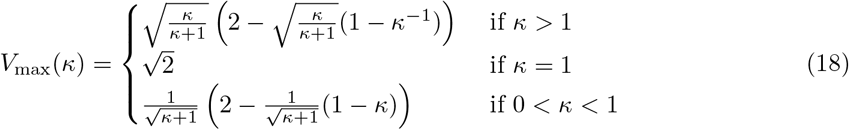
and is plotted in Figure 8. As *κ* → 0 or ∞, the improvement ratio *V*_max_ degrades as one environment becomes favored over the other. In these extreme scenarios, either OA or OB become optimal for increasingly larger regions.

**Figure 7:**
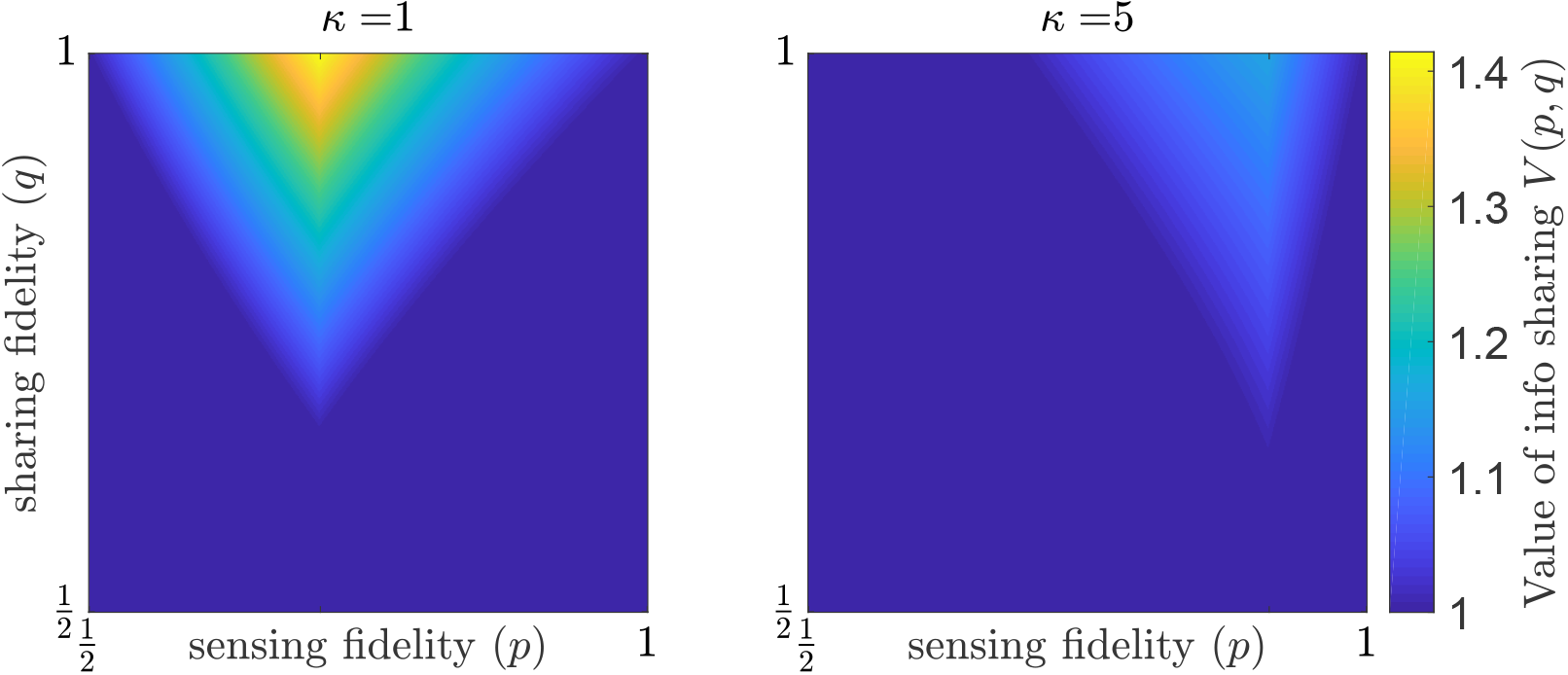
The fitness value of information sharing *V*(*p*, *q*, *κ*). (Left) When *κ* = 1, the maximum value of *V* is attained at (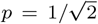,*q* = 1,*κ* = 1), where information sharing leads to approximately a 41% increase in fitness over no sharing. (Right) When *κ* = 5, information sharing leads to approximately 15% increase in fitness at best (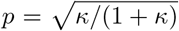, *q* = 1).

**Figure 8.**
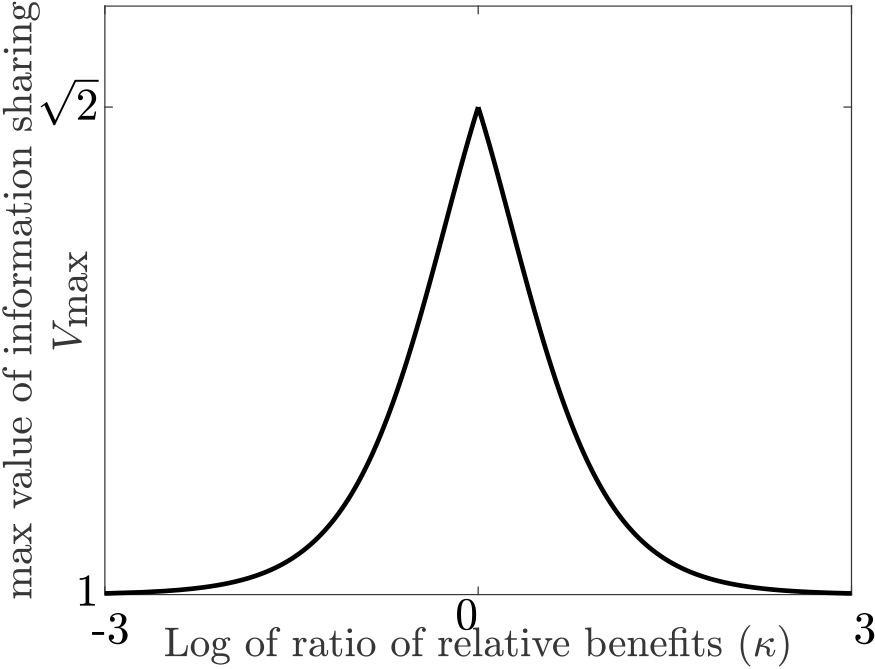
The maximum value of information sharing *V*_max_(*κ*) (Eq. (18)), which is the value of *V*(*p*, *q*, *κ*) attained at *q* = 1 and 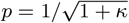 when *κ* > 1. 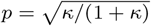 when *κ* ≥ 1. The peak of fitness improvement occurs when no environment is favored over the other (*κ* = 1), where there is an improvement ratio of 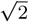. The value for *V*_max_ is *f_*pq*_*(ML_*B*_)/*c*_*B*_ when *κ* < 1, and *f_*pq*_*(ML_*A*_)/*c*_*A*_ when *κ* > 1 (they coincide when *κ* = 1). The fitness improvement ratio degrades as *κ* → 0 or ∞, i.e. when one environment is favored over the other.

We also note that *V*(*p*, *q*, *κ*) increases quadratically in *q* for *q* > *q*_*c*_(*p*). Hence, the rate at which *V* increases with respect to *q*, 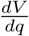, increases linearly in *q*. This is due to a majority logic strategy being the fitness maximizer in this region. We also note that *V*(*p*, *q*, *κ*) decreases as *p* deviates away from the intermediate value 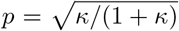. At the extremes *p* =1/2 and *p* =1, the strategies *OA* and FC begin to dominate, respectively, as the line *q*_*c*_ (*p*) tapers to 1.

## 4. Discussion

We have presented a two-player, two action coordination game in a fluctuating environment where players independently sense private cues from the environment and share their cues with each other. In our analysis, we found the optimal strategies that promote coordination across all levels of sensing and sharing fidelities (Figures 4 and 6). When individual sensing is very reliable, there is no need to share signals because players can accurately infer the environmental state independently. When sensing is unreliable, players prefer to ignore their information altogether and always commit to a single action. The “majority logic” strategy is optimal when private information has intermediate fidelity and social information has high fidelity. This strategy highlights the importance of information sharing because they are the only optimal strategies that utilize the shared social cues. Because these optimal strategies are derived from a first-principles approach, their appearance in our model offers insight into the mechanisms that maintain coordinated group behaviors.

The Majority Logic strategies strike a balance between when to use information as a predictor of the environment and when to use information as a means to coordinate. Essentially, it allows players to coordinate when their individual inferences (private information) about the environment conflict, and to choose the correct action when they agree. The interplay of private and social information in our model draws similarities to Condorcet’s jury theorem, where in King and Cowlishaw (2007), states that good reliability of private information is a requirement to effectively make group decisions based on social information in the large population limit. In their model, individuals disregard social information when private information is poor. When it is more accurate, individuals rely more on social information to make a decision. In our model, these two situations are akin to the fitness maximizing regions of OA and OB for poor sensing fidelity, and the region where ML thrives, respectively. These results also corroborate with controlled experimental lab work conducted on nine-spined sticklebacks (van Bergen et al., 2004). When private experience about foraging sites was 100% reliable, sticklebacks based foraging decisions only on private information. When it was less reliable, they followed social cues instead. Without perfect sensing capabilities, organisms need to rely on social information to survive (van Bergen et al., 2004; King and Cowlishaw, 2007; Pérez-Escudero and de Polavieja, 2011; Arganda et al., 2012; Miller et al., 2013).

In the region of unreliable private information and reliable social information (Fig 5, right), there is an abundance of Nash equilibria (ten) that are suboptimal to the OA and OB strategies. This suggests that in the context of coordination games, social cues serve as a coordination device rather than as an additional source of information about the environment. However, the strategies included in this region, which includes majority logic and its variants, will often coordinate on the wrong action. This is because the social cues carry no information about the environment, as the private cues are themselves uninformative. The best an individual can do to infer the environment is simply to guess. However, if both guess independently, they will only coordinate 1/4 of the time. If they play OA, they are guaranteed to coordinate for the fraction of time the environment spends in state *e*_*A*_. Hence, committing to a single action corresponding to the most frequent environment (OA or OB) is the best the group can do when information about the environment is poor. Therefore in principle, information sharing is useful only when the information that is being shared is itself reliable.

Our game-theoretic model portrays situations where communicating individuals must coordinate behaviors in uncertain fluctuating environments. These situations pervade collective behaviors in groups of organisms across the animal kingdom from social insects to bacterial colonies. The information flow in our model is particularly inspired by quorum sensing in bacterial populations. Such a communication system enables bacteria to display complex social behaviors (West et al., 2007). Therefore, game theory is a natural framework in which to study microbial decision-making to consolidate experimental understanding of the phenomena. Advancing such knowledge may also have therapeutic applications. For example, a deeper understanding of how bacteria communicate to form harmful biofilms presents an opportunity to develop methods to inhibit the fidelity of quorum sensing systems (Rutherford and Bassler, 2012; Popat et al., 2015).

Several questions remain in our study. In animal communication, better quality signalling entails increasing fitness costs (Bergstrom and Lachmann, 1997; Brown and Johnstone, 2001; Skyrms, 2010; Meacham et al., 2013; Huttegger et al., 2014). Our work has not yet investigated such evolutionary tradeoffs. Instead, we have presented a systematic, centralized analysis of the optimal strategies given a fixed, costless communication system. Consequently, we have yet to address whether the optimal strategies identified are stable in an evolutionary sense, with or without costly signalling.

Moving forward, several evolutionary dynamics can be applied to our model in the context of population games (Sandholm, 2010). Population games are a framework to describe the interactions between a well-mixed, continuous mass of agents that select from the same set of strategies. In the population, interactions between agents are probabilistic and pair-wise, which allows two-player normal form games to be represented as population games. Such a framework works well with our aim to describe a population of organisms that can adopt a variety of communication-based strategies, *e.g.,* quorum sensing bacteria. Furthermore, due to the potential structure of our games 𝒢_*p*_ and 𝒢_*pq*_ (see Remark 3), they admit evolutionary dynamics that have certain stability guarantees (Sandholm, 2001, 2010). Embedding our model into population games will be necessary to identify which local maximizers of average fitness are likely to be reached. In doing so, we hope that our model encourages the integration of social interactions and communication into efforts to understand coordination, cooperation, and conflict in complex environments.

## Acknowledgements

This work is supported by ARO grant #W911NF-14-1-0402. We thank William H. Sandholm, Georgy Loginov, and William C. Ratcliff for their comments on the manuscript. We thank Marvin Whiteley and Dan Cornforth for insightful discussions. We thank two anonymous reviewers for their constructive feedback. K.P. and C.E. also thank Matthieu Bloch for insightful discussions.

## Code availability

The code for our simulations is available online at http://dx.doi.org/10.5281/zenodo.1179068.

